# Gut Microbiota Dysbiosis and Short-chain Fatty Acid alterations in Pediatric New-Onset Type 1 diabetes with Ketoacidosis

**DOI:** 10.1101/2025.04.22.650027

**Authors:** Yaru Liu, Hu Lin, Mingqiang Zhu, Xuefeng Chen, Zhu Yu, Guanping Dong, Yan Ni, Junfen Fu

## Abstract

**Objective:** Diabetic ketoacidosis (DKA) stands as the most common acute hyperglycaemic complication with a relatively high mortality rate among children diagnosed with type 1 diabetes (T1D). Previous studies have shed light on gut dysbiosis in newly-onset T1D children, but research targeting the gut microbiota and microbial metabolites in patients with DKA remains scarce.

**Methods:** Shotgun metagenomic analysis was conducted on fecal samples from 96 newly diagnosed T1D children, including 32 patients who presented with DKA upon admission. Short-chain fatty acids (SCFAs) were quantified using gas chromatography/mass spectrometry (GC/MS). Comparative analyses were performed on the gut microbiome and SCFA levels between DKA and non-DKA subgroups.

**Results:** Gut microbiota composition differed between DKA and non-DKA groups. While alpha-diversity remained consistent, beta-diversity showed marked differences between the two groups, and the phyla *Firmicutes* and *Bacteroidetes* exhibited predominance in the DKA group. DKA was associated with increased potential pathogens and a significant depletion of SCFA-producing genera (*Anaerobutyricum*, *Dialister*, *Ruminococcus*, *Roseburia*, *Dorea*, and *Butyricicoccus*), accompanied by reduced SCFA levels. Further correlation analysis revealed strong associations between SCFAs, SCFA-producing microbiota, and disease severity indicators. Propionic acid and butyric acid mediated the association between SCFA-producing bacteria and DKA.

**Conclusion:** This study first reported reductions in SCFAs and SCFA-producing bacteria in the fecal samples of T1D children at DKA onset. Our findings highlight SCFAs and their producers as protective biomarkers of DKA and suggest that targeting gut microbiota and SCFA metabolism may offer a potential therapeutic strategy for DKA.

**Importance:** While gut microbiota imbalances have been linked to type 1 diabetes (T1D), little is known about the role of gut microbes and their metabolites during acute diabetic ketoacidosis (DKA), an acute and life-threatening complication. This study examined the gut microbiome and levels of short-chain fatty acids (SCFAs) in fecal samples from children with new-onset T1D with and without DKA. DKA was characterized by distinct microbial signatures, including increased potential pathogens, a loss of beneficial SCFA-producing bacteria, and consistently lower SCFA levels. These changes closely correlated with indicators of disease severity, and further mediation analysis revealed the role of SCFAs in linking gut microbiota to DKA. These findings suggest that SCFAs and their producers may serve as protective markers for DKA, highlighting the potential of microbiome-based interventions to improve outcomes in children facing this critical complication.

## Introduction

Type 1 diabetes (T1D) is an autoimmune disease characterized by insufficient endogenous insulin due to immune damage to pancreatic islet β cells, resulting from heterogeneous genetic and environmental factors ^1^. T1D can occur at any age but typically develops in children and adolescents ^2^. The global incidence of T1D is estimated to be approximately 15/100,000 ^3^, which has been continuously increasing over the past years. Multiple regional studies in China have also shown a rising incidence of T1D in children ^4–6^. Lacking a complete cure, children diagnosed with T1D require lifelong use of insulin and blood glucose monitoring. Poor compliance with insulin injections contributes to a higher incidence of diabetes-related complications, which poses a heavy disease burden ^7^. The most common acute complication, diabetic ketoacidosis (DKA), is the leading cause of death in children with T1D ^8^. Previous studies from developing countries have reported a notably high mortality rate among children with DKA, ranging from 3% to 13% ^9^. However, research on physiological changes occurring during DKA remains limited.

As the body’s largest immune organ, the gut hosts a complex microbiota ecosystem essential for preserving the integrity of the intestinal mucosal barrier. Gut microbiota dysbiosis has been proven to be associated with various autoimmune diseases, including T1D ^10^. Several studies have reported a decreased gut microbiota diversity in children with T1D compared to non-diabetic controls ^11,12^. Accordingly, beneficial gut microbial metabolites also decreased, such as short-chain fatty acids (SCFAs), which maintain the immune tolerance by promoting CD4^+^ regulatory T (Treg) cells and suppressing autoreactive T cells in the gut ^13^. Compared to healthy controls, various SCFAs in the feces of children with T1D were lowered ^12^. The Environmental Determinants of Diabetes in the Young (TEDDY) study, conducted in the United States, Sweden, Finland, and other countries, also identified a reduced abundance of SCFA-producing bacterial pathways in children with islet autoantibodies or T1D ^14^. Growing evidence suggests that maintaining gut structure and function may serve as a potential therapeutic approach for T1D. Fecal microbiota transplantation (FMT) has shown potential for alleviating newly diagnosed T1D in humans or animals ^15,16^, while probiotics (e.g., *Bifidobacterium*, *Lactobacillus*) and maternal SCFA supplementation can mitigate T1D risk ^17–19^.

Previous research has primarily focused on the gut microbiota and metabolites in newly diagnosed T1D, while studies addressing the acute complication, DKA, remain scarce.

Based on the clinical dataset alongside fecal metagenomics and metabolomics, this study investigated the alterations in gut microbiota and SCFA levels in children with T1D experiencing DKA and assessed their correlations with disease indicators. Our study aims to explore potential biomarkers and new therapeutic strategies for T1D and DKA management.

## Methods

### Study population

The study was conducted following ethical approval granted by the Ethics Committee of the Children’s Hospital, Zhejiang University School of Medicine (Ethics Approval Number: 2020-IRB-081). The study cohort comprised newly diagnosed children and adolescents with T1D, recruited from the Department of Endocrinology at the Children’s Hospital, Zhejiang University School of Medicine, between May 2016 and June 2023. The exclusion criteria for the T1D group were as follows: (1) individuals diagnosed with type 2 diabetes, mixed diabetes, monogenic diabetes confirmed through genetic testing, and other specific types of diabetes; (2) children or their parents who declined to participate in the study. From all participants, we obtained written informed consent before enrollment. The following research workflow is shown in **Figure 1**.

**Figure 1.**
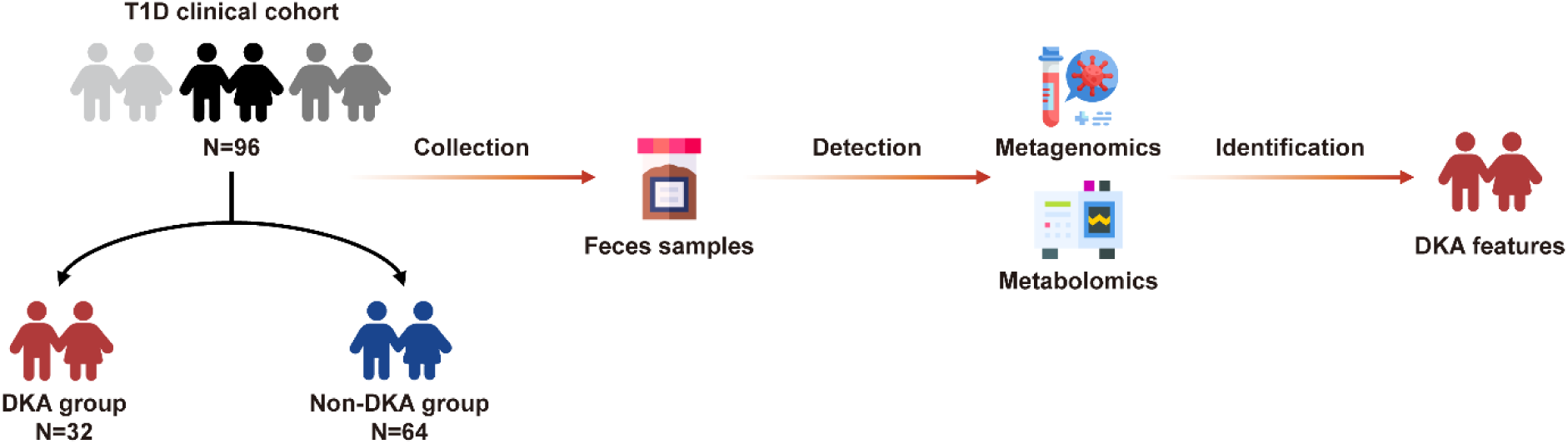
Research flowchart for T1D subjects. Shotgun metagenomic analysis was performed on fecal samples obtained from 96 newly diagnosed T1D children, including 32 with DKA upon admission. Short-chain fatty acids (SCFAs) were quantified using gas chromatography/mass spectrometry (GC/MS). Comparative and correlation analyses were conducted on the gut microbiome, SCFA levels, and clinical features between DKA and non-DKA groups.

### Clinical data collection

At baseline, we collected demographic and clinical data of newly diagnosed T1D patients. Body mass index (BMI) and BMI z-score were calculated to assess growth and nutritional status. Comprehensive laboratory evaluations were performed, including routine blood and urine tests, blood glucose levels, serum insulin, C-peptide, glycated hemoglobin (HbA1c), diabetes-related autoantibodies, lipid profiles, liver and kidney function tests, thyroid function, blood gas analysis, electrolyte levels, and urinary microalbumin measurements.

### Sample collection and DNA extraction

Fresh fecal samples from newly diagnosed T1D patients were collected in disposable sterile containers at baseline and stored at -80°C. Genomic DNA (∼0.2 µg) was extracted for library preparation. DNA fragments were sonicated to an average length of 350 bp, end-repaired, A-tailed, and PCR-amplified. The PCR products were purified using the AMPure XP system (Beverly, USA), quality-controlled with the Agilent 5400 system (Agilent, USA), and quantified by qPCR (1.5 nM). Libraries meeting quality standards were pooled and sequenced on the Illumina platform (PE150 strategy) by Novogene Bioinformatics Technology Co., Ltd, Beijing, China. Raw fluorescent images were converted to sequence data in FASTQ format. Raw data quality was assessed with FastQC (v0.11.9), and Trimmomatic (v0.39) was employed to filter out adapters, duplicates, and low-quality reads. Clean reads were aligned to the host genome using BWA (0.7.17-r1188) to remove high-similarity contaminant sequences.

### Metagenomic sequencing

Clean reads were assembled using MEGAHIT (v1.2.9) and assessed with QUAST (v5.2.0). ORFs in the contigs were predicted with MetaGeneMark (v3.38), filtered (≥100 nt), and translated into amino acid sequences. Non-redundant gene sets were constructed using CD-HIT (v4.8.1) (identity 95%, coverage 90%), and clean reads were mapped using BWA. Taxonomic annotation was performed by aligning gene sets against the NCBI NR database with DIAMOND (v2.1.6.160), classifying species from phylum to species level. Functional profiling was conducted via DIAMOND against protein functional databases, integrating gene abundance. Microbial diversity was analyzed in R software (v4.3.2) using the Vegan package (Shannon and Simpson indices for alpha diversity, PCoA with Bray-Curtis distances for beta diversity). Differential abundance analysis was performed with the DESeq package, while multivariate analysis used partial least squares discriminant analysis (PLS-DA).

### Short-chain fatty acid detection

Fecal samples were centrifuged, and 200 μL of supernatant was mixed with 100 μL of 15% phosphoric acid, 20 μL of an internal standard (4-methylvaleric acid, 375 μg/mL), and 280μL of ether. After homogenization and centrifugation, the supernatant was analyzed using a Thermo Trace 1310 gas chromatography system coupled with a Thermo ISQ LT mass spectrometer (Thermo Fisher Scientific, USA). Metabolite concentrations were quantified via standard curves, determining the content of acetic acid (AA), propionic acid (PA), butyric acid (BA), valeric acid (VA), caproic acid (CA), isobutyric acid (IsoBA), and isovaleric acid (IsoVA).

### Statistical analysis

Statistical analyses were performed using R software (v4.3.2). For continuous variables, normally distributed data were compared using an independent samples t-test, while non-normally distributed data were analyzed by the Mann-Whitney U test. Categorical variables were compared using either chi-square or Fisher’s exact test, as appropriate. The Kruskal-Wallis rank sum test was employed to assess differences in species abundance across groups. Spearman correlation analysis was conducted to explore correlations between differential species, metabolites, and clinical indicators. We further utilized Gephi software (v0.10.1) to construct the correlation network of the gut microbiota. For mediation analysis, missing values in SCFA contents were imputed using the mean values. To estimate the mediation effects of SCFAs in the relationship between SCFA-producing microbiota and DKA, we employed the mediation package (v4.5.0) and mediation function from the bruceR package (v2024.6). Mediation proportion and 95% confidence interval (CI) for individual SCFA in the association between specific microbes and DKA were estimated using the non-parametric bootstrap method with 5000 resamples. A *P*-value < 0.05 was considered statistically significant.

## Results

### Clinical features

This study included 96 children with newly diagnosed T1D, of whom 32 presented with DKA upon admission, while the remaining 64 were assigned to the non-DKA group (**Figure 1**). Age and sex were comparable between the two groups (**Table 1**). A comparison of demographic data showed lower BMI and BMI z-score in the DKA group, along with higher systolic blood pressure (SBP) compared to the non-DKA group. Metabolic disorders in the DKA group included elevated blood glucose, acidosis, and lower pH, bicarbonate, and actual base excess (ABE). Serum lipid tests prompted possible dyslipidemia in the DKA children, characterized by higher levels of low-density lipoprotein cholesterol (LDL-c), Apolipoprotein B, triglycerides (TG), and total cholesterols, with lower levels of high-density lipoprotein cholesterol (HDL-c), Apolipoprotein A, and bile acids. Thyroid hormones were significantly reduced in the DKA group, consistent with previously reported non-thyroidal illness syndrome (NTI) or euthyroid sick syndrome (ESS) ^20,21^. Retinol-binding protein (RBP) and complement component 3 (C3) levels were decreased, whereas interleukin 10 (IL10) was elevated during DKA onset. Urinary tests showed higher levels of proteins, ketones, and glucose in DKA children. At discharge, insulin use per weight and insulin dose-adjusted glycated hemoglobin A1c (IDAA1c) were higher in the DKA group, indicating a more severe disease phenotype.

**Table 1.** Clinical characteristics of T1D subjects. BMI, body mass index; SBP, systolic blood pressure; DBP, diastolic blood pressure; HbA1c, glycated hemoglobin; ABE, actual base excess; HDL-c, high-density lipoprotein cholesterol; LDL-c, low-density lipoprotein cholesterol; TG, triglycerides; ALT, alanine transaminase; AST, aspartate aminotransferase; WBC, white blood cells; TSH, thyroid-stimulating hormone; T3, triiodothyronine; T4, thyroxine; fT3, free triiodothyronine; fT4, free thyroxine; IGF1, insulin-like growth factor 1; IGFBP3, insulin-like growth factor binding protein 3; RBP, retinol-binding protein; C3, complement component 3; IL10, interleukin 10; CD3, CD3 positive T cells; CD4, CD4 positive T cells; CD8, CD8 positive T cells; Urine pr/cr, urine protein to creatinine; IDAA1c, insulin dose-adjusted glycated hemoglobin A1c. * *P* < 0.05, ** *P* < 0.01, *** *P* < 0.001.

| Characteristics | DKA (n=32, 33.3%) | Non-DKA (n=64, 66.7%) | *P*-value |
| --- | --- | --- | --- |
| Demographic data |  |  |  |
| Male | 13 (40.6%) | 27 (42.4%) | 1.000 |
| Age (years) | 6.21 (4.06, 10.28) | 7.11 (5.57, 10.13) | 0.419 |
| Weight (kg) | 20.25 (14.5, 24.75) | 22.75 (18.38, 28.63) | 0.074 |
| BMI (kg/m<sup>2</sup>) | 13.98 (13.22, 15.31) | 14.94 (13.81, 16.51) | 0.041* |
| BMI z-score | -1.19 (-1.85, -0.81) | -0.49 (-1.01, 0.14) | 0.008** |
| SBP (mmHg) | 113.38±12.83 | 107.95±10.29 | 0.043* |
| DBP (mmHg) | 71.09±12.67 | 68.86±8.68 | 0.374 |
| Blood sugar and gas analysis |  |  |  |
| HbA1c (%) | 13.18±1.78 | 12.71±2.13 | 0.301 |
| C-peptide<br>(pmol/L) | 86 (37, 106.5) | 94 (52, 174) | 0.157 |
| pH | 7.22 (7.13, 7.28) | 7.37 (7.35, 7.41) | 0.000*** |
| Glucose (mmol/L) | 21.96±7.65 | 16.66±9.25 | 0.004** |
| Chlorine (mmol/L) | 112.5 (107, 116.75) | 105 (103, 109) | 0.000*** |
| Calcium (mmol/L) | 1.32 (1.25, 1.36) | 1.19 (1.15, 1.23) | 0.000*** |
| Bicarbonate | 11 (5.88, 14.2) | 22 (19.4, 24.45) | 0.000*** |
| (mmol/L) |  |  |  |
| ABE (mmol/L) | -16.2 (-22.6, -10.55) | -2.4 (-4.9, -0.73) | 0.000*** |
| Serum lipid test |  |  |  |
| HDL-c (mmol/L) | 1.09 (0.87, 1.27) | 1.26 (0.98, 1.43) | 0.031* |
| LDL-c (mmol/L) | 3.33 (2.95, 3.93) | 2.96 (2.14, 3.31) | 0.010* |
| Apolipoprotein A1 | 1.12 (0.91, 1.3) | 1.3 (1.02, 1.48) | 0.027* |
| (g/L) |  |  |  |
| Apolipoprotein B | 1.06 (0.96, 1.2) | 0.81 (0.64, 1.03) | 0.000*** |
| (g/L) |  |  |  |
| Lipoprotein A | 24 (17, 59) | 84 (37, 171.5) | 0.000*** |
| (mg/L) |  |  |  |
| TG (mmol/L) | 1.72 (1.12, 3.58) | 0.98 (0.66, 1.29) | 0.000*** |
| Cholesterols | 5.07 (4.59, 6.32) | 4.73 (3.76, 5.5) | 0.011* |
| (mmol/L) |  |  |  |
| Bile acids | 2 (1.45, 3.15) | 4.5 (2.65, 7.88) | 0.000*** |
| (μmol/L) |  |  |  |
| ALT (U/L) | 12 (10.25, 15) | 14 (11, 17.75) | 0.039* |
| AST (U/L) | 20 (15, 25) | 25 (22, 30.5) | 0.002** |
| Diabetic autoantibody |  |  |  |
| Autoantibody+ | 26 (81.2%) | 41 (64.1%) | 0.163 |
| Blood routine test |  |  |  |
| WBC ($\times 10^9/L$) | 10.38 (8.04, 17.23) | 6.69 (5.5, 8.31) | 0.000*** |
| Neutrophils/WBC | 64.37±16.57 | 46.08±16.79 | 0.000*** |
| (%) |  |  |  |
| Haemoglobin | 133.74±13.42 | 128.77±10.47 | 0.098 |
| (g/L) |  |  |  |
| Platelets (×10<sup>9</sup>/L) | 300.48±89.67 | 262.7±65.19 | 0.057 |
| Endocrine hormone levels |  |  |  |
| TSH (mIU/L) | 0.78 (0.39, 1.3) | 1.95 (1.2, 2.74) | 0.000*** |
| T3 (nmol/L) | 0.54 (0.39, 0.73) | 1.05 (0.83, 1.24) | 0.000*** |
| T4 (nmol/L) | 58.64±21.24 | 86.45±19.93 | 0.000*** |
| fT3 (pmol/L) | 2.14 (1.82, 2.71) | 3.56 (3.11, 4.22) | 0.000*** |
| fT4 (mIU/L) | 9.69±2.39 | 12.69±2.23 | 0.000*** |
| Cortisol (µg/dL) | 19.1 (17.1, 29.5) | 16.15 (11.88, 19.48) | 0.001** |
| IGF1 (ng/mL) | 41.9 (32, 66.7) | 71.6 (61.9, 145.5) | 0.007** |
| IGFBP3 (µg/mL) | 1.86 (1.5, 2.27) | 2.89 (2.01, 3.97) | 0.028* |
| Immune tests |  |  |  |
| RBP (mg/L) | 17.5 (13.7, 23.1) | 21.3 (17.25, 26.8) | 0.005** |
| C3 (g/L) | 0.8±0.19 | 0.9±0.19 | 0.018* |
| IL10 (pg/mL) | 5.5 (3.38, 9) | 3.65 (2.85, 4.65) | 0.009** |
| CD3 (%) | 67.2 (64.35, 71.45) | 73.8 (68.36, 77.23) | 0.012* |
| CD4 (%) | 35.2±6.89 | 38.67±6.23 | 0.033* |
| CD4/CD8 | 1.35 (1.19, 1.61) | 1.54 (1.2, 1.83) | 0.280 |
| Urine tests |  |  |  |
| Urine pr/cr | 0.63 (0.3, 1.47) | 0.21 (0.13, 0.29) | 0.000*** |
| (mg/mgCr) |  |  |  |
| Urine | 350.75 (171.18, 647.85) | 91.5 (62.28, 159.05) | 0.000*** |
| microprotein (mg) |  |  |  |
| Urine ketones + | 18 (56.3%) | 32 (50.0%) | 0.000*** |
| Urine glucose + | 29 (90.6%) | 49 (76.7%) | 0.000*** |
| Therapeutic assessment |  |  |  |
| Insulin per weight<br>(U/kg/d) | 0.9±0.24 | 0.73±0.29 | 0.004** |
| IDAA1c | 16.7±1.95 | 15.62±2.74 | 0.048* |
| Insulin pump use | 13 (40.6%) | 29 (45.3%) | 0.930 |

### Gut dysbiosis with an imbalance of pathogens and symbiotics in DKA

The gut microbiota composition between DKA and non-DKA groups was similar, as shown in the relative abundance plot (**Figure 2A**). The top four phyla in both groups were *Firmicutes* (also known as *Bacillota*), *Bacteroidetes* (or *Bacteroidota*), *Actinobacteria* (or *Actinomycetota*), and *Proteobacteria* (or *Pseudomonadota*). However, compared to the non-DKA group, the relative abundances of phylum *Firmicutes* and *Bacteroidetes* were lower in the DKA group (**Figure 2B**). Although alpha-diversity remained consistent between DKA and non-DKA subjects, beta-diversity revealed pronounced differences between the two groups (*P* < 0.05) (**Figure 2C, D** and **Supplementary Figure 1A**). Using PLS-DA for multivariate analysis, DKA and non-DKA groups were distinguished at phylum, genus, and species levels (**Figure 2D** and **Supplementary Figure 1B**). These findings suggest distinct microbial composition between newly diagnosed children with and without DKA.

**Figure 2.**
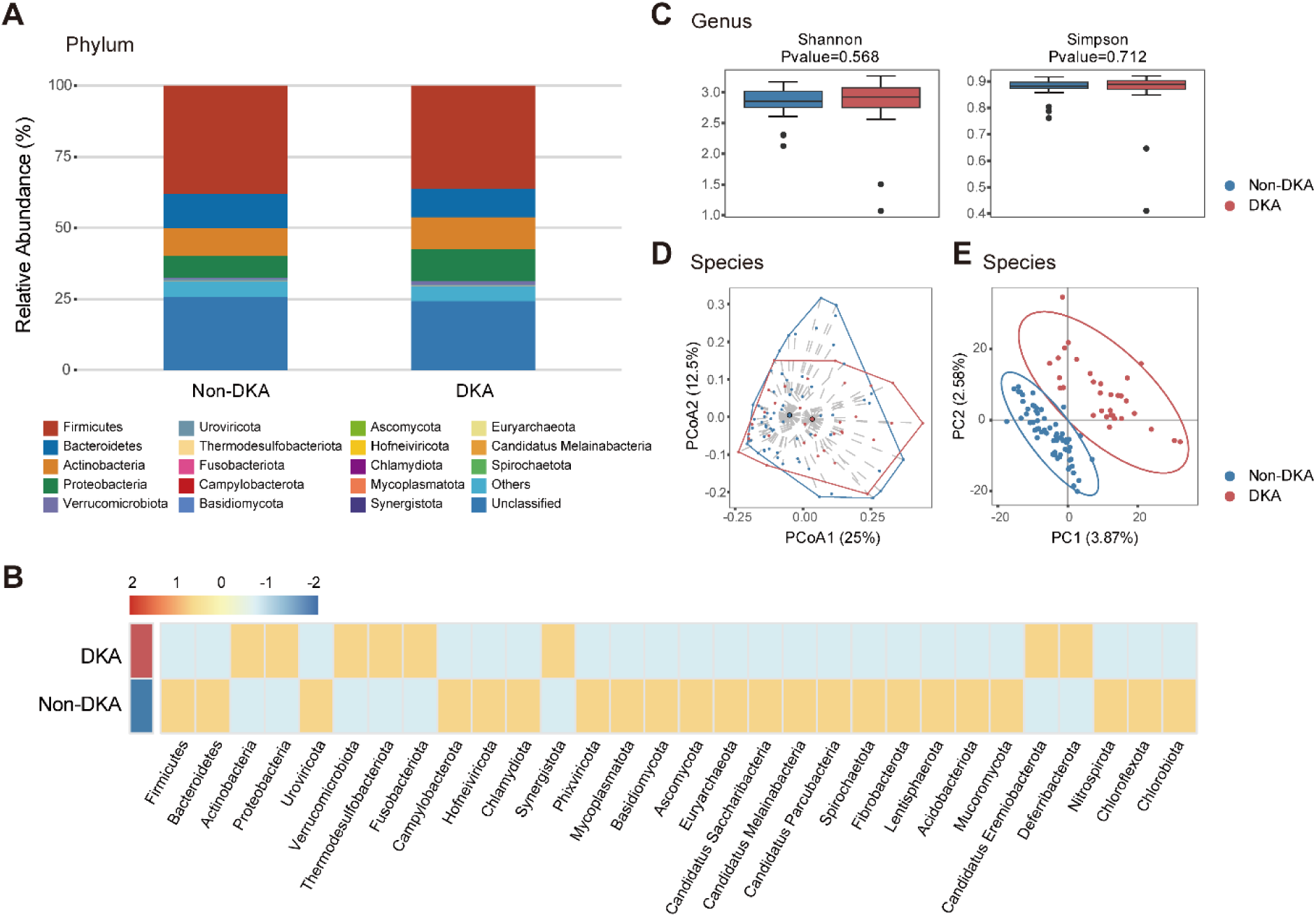
Gut microbiota differences between DKA and non-DKA groups. (**A**) Gut microbiota composition at the phylum level showed similarities between the DKA and non-DKA groups. (**B**) Z-score heatmap revealed lower relative abundances of the phyla *Firmicutes* and *Bacteroidetes* in the DKA group. (**C**) Alpha-diversity was consistent between the two groups. (**D**) Principal coordinates analysis (PCoA) showed significant beta-diversity differences at the species level (*P* = 0.010). (**E**) Through partial least squares discriminant analysis (PLS-DA), DKA and non-DKA groups were distinguished at the species level (*P* < 0.05).

After a literature review, we classified the gut microbiota into possible pathogens and potential symbiotics and compared their differences in abundance between the two groups. Potential pathogenic microorganisms were enriched in the DKA group, with increased abundances of *Erysipelotrichi*, *Alphaproteobacteria*, and *Mucoromycetes* at the class level; *Erysipelotrichale*, *Mucorales*, and *Mycobacteriales* at the order level; and *Staphylococcaceae*, *Nitrobacteraceae*, *Coprobacillaceae*, *Burkholderiaceae*, and *Enterococcaceae* at the family level (**Supplementary Figure 2A**). Conversely, symbiotic bacteria with potential host benefits were reduced in the DKA group, including the order *Methylococcales*, the family *Methylococcaceae* and *Oscillospiraceae*, and the genus *Anaerobutyricum*, *Dialister*, *Ruminococcus*, *Roseburia*, *Dorea*, and *Butyricicoccus* (**Supplementary Figure 2B**). We constructed network diagrams to illustrate the correlations among microbiota significantly increased and decreased in the DKA group, respectively. More increased microbes in the DKA group were possible pathogens (**Figure 3A**), while decreased microbes were primarily potential symbiotics (**Figure 3B**). Notably, significant positive correlations were observed among the decreasing microbes, especially the potential symbiotics, prompting their close symbiotic effects. These findings highlighted gut microbiota dysbiosis associated with DKA, characterized by increased potential pathogens and decreased beneficial symbiotics.

**Figure 3.**
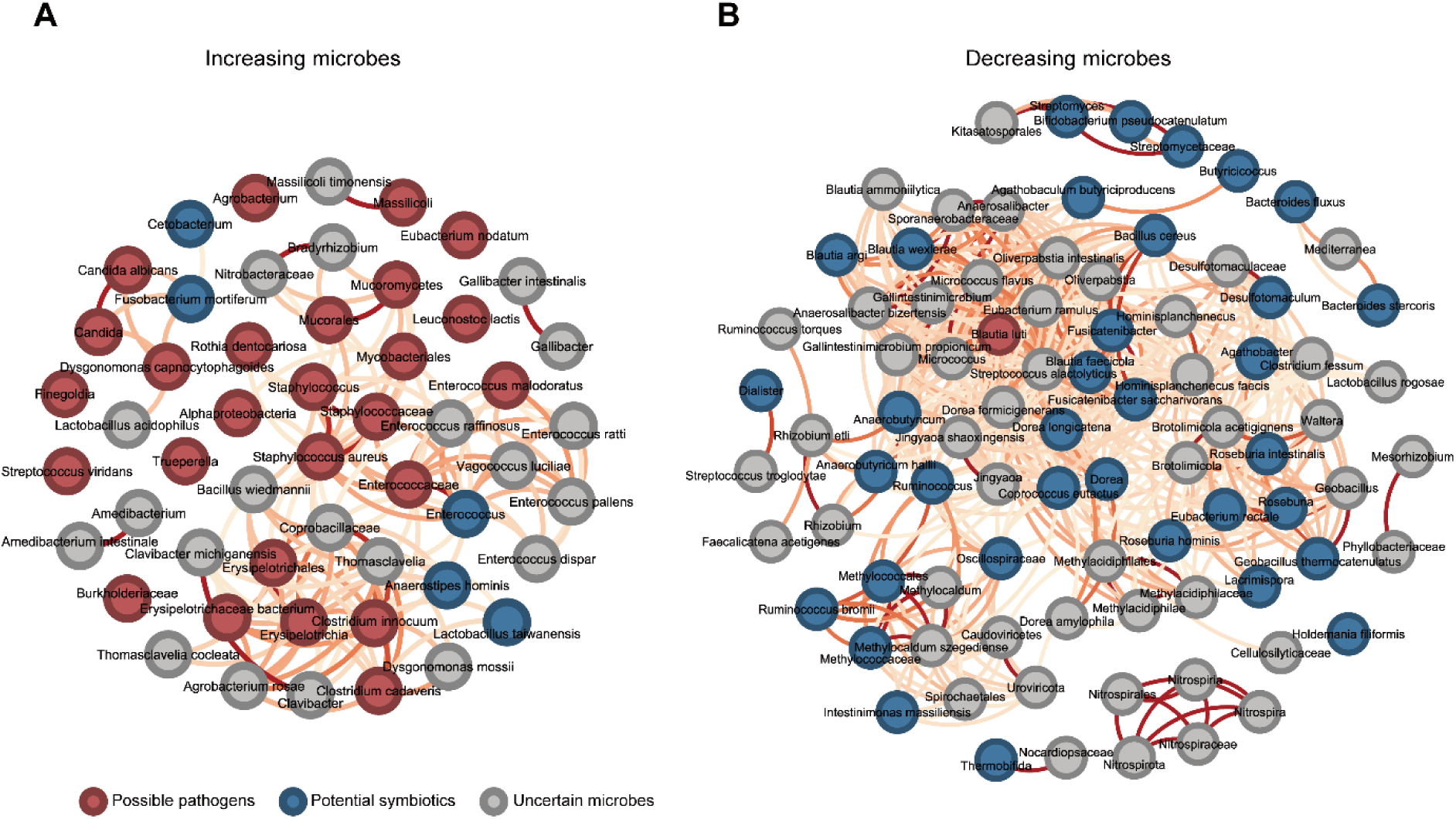
Gut microbiota network revealing an imbalance of pathogens and symbiotics in the DKA group. Correlation networks of increasing (**A**) and decreasing (**B**) microbes at all microbial levels in the DKA group are generated using Gephi software (v0.10.1). Only correlations with absolute coefficients ≥ 0.5 and a *P*-value ≥ 0.05 are displayed. All correlations were positive, represented by red lines, with intensity reflecting the strength of the correlation coefficients.

Additionally, we observed changes in the abundance of several beneficial microbes capable of producing SCFAs (**Figure 4**). At the genus level, *Anaerobutyricum*, *Dialister*, *Ruminococcus*, *Dorea*, *Thermobifida*, *Agathobacter*, *Butyricicoccus*, and *Fusicanibacter* exhibited significantly lower abundances in the DKA group compared to the non-DKA group. At the species level, *Anaerobutyricum hallii*, *Roseburia hominis*, Roseburia intestinalis, Intestinimonas massiliensis, *Fusicanibacter saccharivorans*, *Agathobaculum butyriciproducens*, *Ruminococcus bromii*, *Eubacterium rectale*, and *Bacteroides stercoris* were more abundant in the non-DKA group. It indicated a notable shift in SCFA-producing bacteria in T1D children at DKA onset.

**Figure 4.**
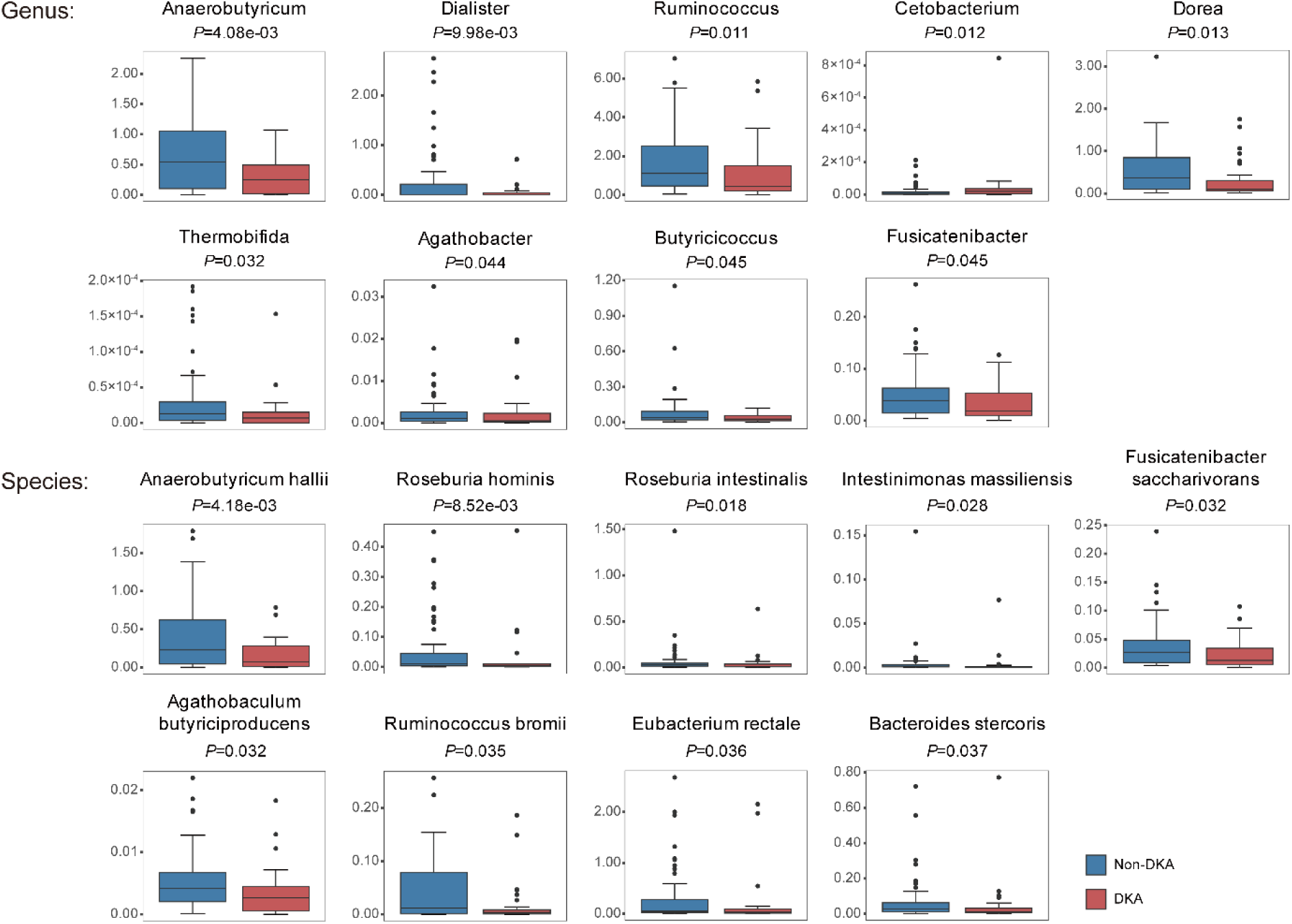
Decreased abundance of SCFA-producing bacteria in the DKA group. Differential analysis identified seventeen SCFA-producing microbes at the phylum and species levels with increased abundance in the DKA group, except for *Cetobacterium*, which exhibited a reduction.

### Decreased levels of Short-chain fatty acids in DKA

Metagenomic analysis revealed a significantly lower abundance of various SCFA-producing gut microbiota in the DKA group compared to the non-DKA group. Consequently, we further assessed and compared the baseline fecal SCFA levels in 76 children with T1D, among which 28 cases (36.8%) developed DKA at disease onset (**Figure 5**). Compared to the non-DKA groups, the levels of six SCFAs, including acetic acid (AA), propionic acid (PA), butyric acid (BA), valeric acid (VA), isobutyric acid (isoBA), and isovaleric acid (isoVA), were significantly lower in the DKA group. This aligned with the metagenomic analysis, suggesting that both SCFA-producing gut microbiota abundance and SCFA levels were reduced during DKA onset. Therefore, SCFAs and SCFA-producing microbiota may be potential biomarkers for DKA.

**Figure 5.**
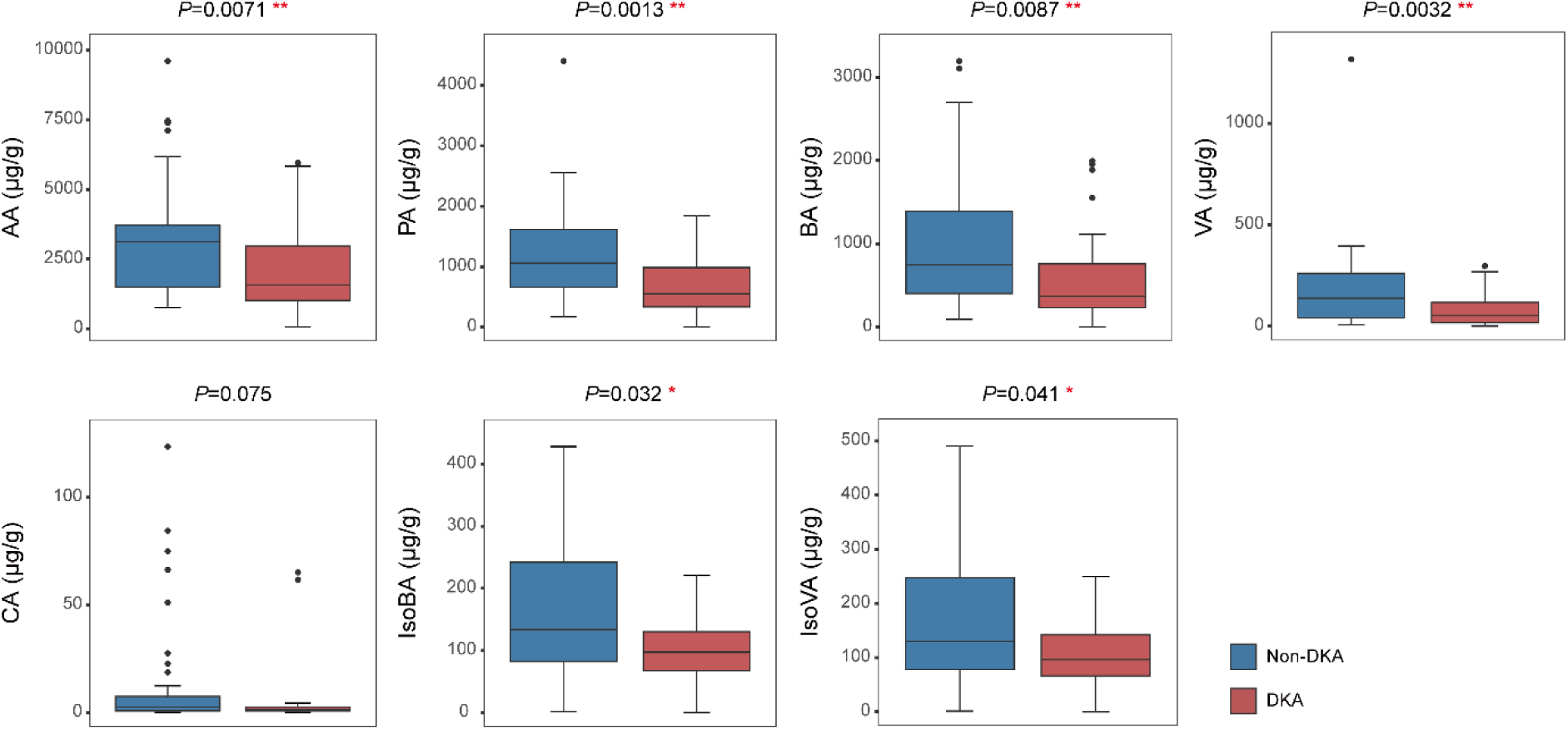
Fecal SCFA levels in the DKA and non-DKA groups. The fecal concentrations of acetic acid (AA), propionic acid (PA), butyric acid (BA), valeric acid (VA), caproic acid (CA), isobutyric acid (isoBA), and isovaleric acid (isoVA) were measured and compared between DKA and non-DKA children. * *P* < 0.05, ** *P* < 0.01.

### Correlations between SCFAs and key bacteria with clinical markers

Next, we performed the Spearman correlations between clinical phenotypes, SCFA-producing bacteria, and SCFAs to elucidate the key features of DKA (**Figure 6A** and **B**). DKA was negatively correlated with various SCFAs, while indicators of acidosis (pH, bicarbonate, and ABE) were positively correlated with SCFAs, suggesting that the severity of acidosis is associated with a decrease in SCFA levels. Blood glucose, HbA1c, and serum C-peptide levels negatively correlated with SCFAs, with only HbA1c and propionic acid exhibiting a significant correlation. Urine glucose and ketones were negatively correlated with propionic acid, indicating that propionic acid may be a marker of DKA severity. Lipid tests, including triglycerides, cholesterol, and Apolipoprotein B, also exhibited negative trends with SCFAs. Apolipoprotein A1, a protective factor against dyslipidemia, was positively correlated with propionic acid, hexanoic acid, and other SCFAs. At the genus level, SCFA-producing microbiota (excluding *Cetobacterium*), were negatively correlated with DKA (**Figure 6A**). Several SCFA-producing genera were positively correlated with acidosis indicators. The genus *Thermoactinomyces* and the species *Intestinimonas massiliensis* and *Ruminococcus bromii*, were negatively correlated with blood glucose indicators (**Figure 6B**). *Butyricicoccus* showed a negative correlation with urine ketones. Particularly in SCFA-producing microbiota, *Roseburia* was negatively correlated with triglycerides and cholesterol. These microbes exhibited significant positive correlations with multiple SCFAs. Thus, SCFAs and their producing bacteria appear as protective factors against DKA.

**Figure 6.**
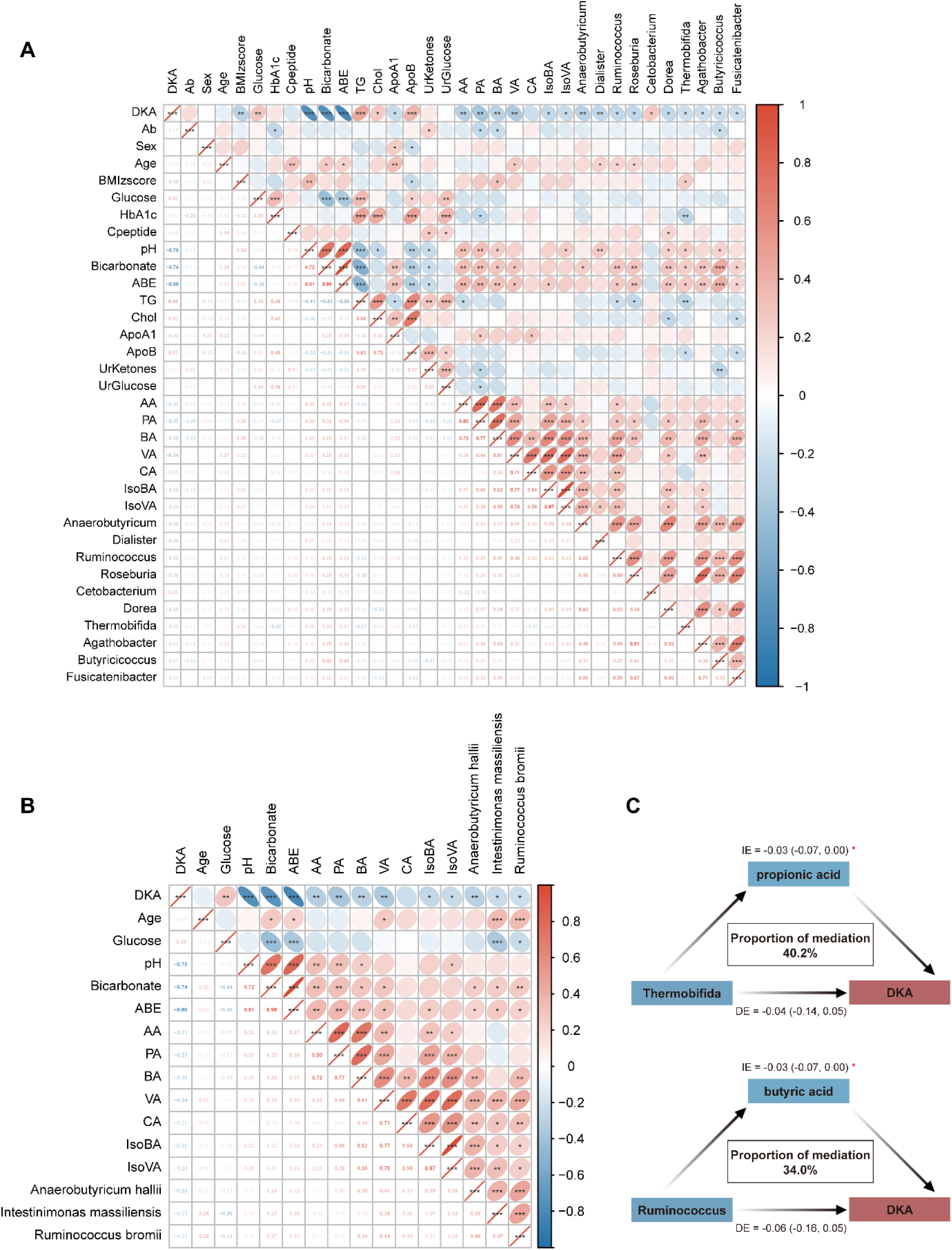
Correlation and mediation analysis of SCFAs, their producing bacteria, and core clinical features of DKA. (**A**) Spearman correlation analysis at phylum level. (**B**) Spearman correlation analysis at species level. The red color indicated positive correlations, while blue denoted negative correlations. The numerical values in the lower left corner of each cell represented the correlation coefficients, and the color intensity in the upper right ellipses reflected the strength of these correlations. (**C**) Mediation analysis of propionic acid and butyric acid in the relationship between *Thermobifida*, *Ruminococcus*, and DKA. IE, indirect effect; DE, direct effect. * *P* < 0.05, ** *P* < 0.01, *** *P* < 0.001.

Additionally, we conducted a mediation analysis to assess whether SCFAs mediated the relationship between SCFA-producing microbiota and DKA (**Supplementary Table 1**). Only PA and BA were significantly involved in the mediation. Specifically, PA mediated the relationship between *Thermobifida* and DKA, accounting for 40.2% of the relationship, and BA mediated the association between *Ruminococcus* and DKA with a contributing proportion of 34.0% (**Figure 6C**). Therefore, SCFAs and their producing bacteria are associated with metabolic dysregulation in DKA, potentially to be protective biomarkers for DKA.

## Discussion

Our study examined the alterations in gut microbiota composition and its microbial product, SCFAs, during the acute complication of DKA in children with T1D. Using metagenomic sequencing and metabolomic analysis, we identified significant differences in gut microbiota between DKA and non-DKA children. This is the first study to report a reduction in SCFAs and their producing bacteria in Chinese children with T1D who developed DKA. Propionic acid, butyric acid, and their producing microbiota, such as the genus *Ruminococcus*, are potential biomarkers for DKA.

SCFAs are metabolic byproducts formed by bacterial fermentation of indigestible polysaccharides in the human gut, primarily by colonic bacteria. Structurally, SCFAs are saturated fatty acids containing 1-6 carbon atoms, including formate, acetate (or acetic acid), propionate (propionic acid), butyrate (butyric acid), valerate (valeric acid), and hexanoate (caproic acid), with acetate, propionate, and butyrate being the most abundant in the human body ^22,23^. Additionally, we measured two branched-chain SCFAs, isobutyrate and isovalerate, which are also microbial metabolites but are present in lower concentrations compared to the primary SCFAs ^24^. After bacterial fermentation in the colon, SCFAs enter the liver via the portal vein and are distributed to target organs, including adipose tissue, immune organs, the brain, pancreas, heart, and kidneys, where they regulate glucose and lipid metabolism and exert anti-inflammatory effects by binding to G-protein coupled receptor 43 (GPR43) ^25^. In the gut, they stimulate the secretion of gut hormones peptide YY (PYY) and glucagon-like peptide-1 (GLP-1) and enhance insulin sensitivity. In the pancreas, SCFAs promote insulin secretion, contributing to glucose regulation ^26^. Butyrate activation of GPR109A receptors in adipocytes inhibits excessive lipolysis and reduces free fatty acid release ^26^. In immune cells, SCFAs modulate leukocyte migration and cytokine release during inflammation. For example, butyrate enhances Treg cell differentiation via GPR109A, exhibiting anti-inflammatory properties ^26,27^. These findings highlight the potential protective role of SCFA supplementation in mitigating acute metabolic disturbances during DKA onset.

In children with DKA, several reduced SCFA-producing genera belong to the *Firmicutes* phylum, *Clostridia* class, and *Eubacteriales* order, including *Anaerobutyricum*, *Ruminococcus*, *Roseburia*, *Dorea*, *Agathobacter*, *Butyricicoccus*, and *Fusicatenibacter*. Except for *Ruminococcus* (*Oscillospiraceae* family) and *Butyricicoccus* (*Clostridiaceae* family), all other genera are classified within the *Lachnospiraceae* family. Many genera within the *Lachnospiraceae* family are primary butyrate-producers, exhibiting superior anti-inflammatory effects to acetate and propionate ^28,29^. For example, *Anaerobutyricum* and *Roseburia* convert carbohydrates to butyrate via the butyryl-CoA: acetate CoA-transferase route ^29,30^. *Roseburia*, a key gut symbiont, constitutes 3 to 15 % of fecal bacteria in healthy individuals and can also produce propionate from deoxyhexose through the propanediol pathway ^31,32^. *Dorea*, another member of the *Lachnospiraceae* family, primarily generates acetate, which serves as a precursor for butyrate synthesis via the butyryl-CoA: acetate CoA-transferase route ^33,34^. *Ruminococcus* (*Oscillospiraceae* family) generates acetate, propionate, and butyrate and has been linked to various autoimmune and metabolic diseases, such as obesity and type 2 diabetes; however, its precise role remains uncertain ^29,35–38^. This study observed a significant reduction in *Ruminococcus* abundance in DKA children, suggesting its protective role in DKA. Notably, butyrate significantly mediated the association between *Ruminococcus* and DKA, highlighting its potential role in disease progression. In contrast, *Cetobacterium* within the *Fusobacteriota* phylum, enriched in DKA, can produce acetate, propionate, and butyrate but is more prevalent in fish intestines than humans ^39,40^. Our study identified reduced abundances of key SCFA-producing bacteria in DKA children, suggesting potential probiotic roles in disease prevention. Additionally, comparative analysis across multiple taxonomic levels revealed a higher abundance of pathogenic bacteria in children with DKA, such as *Erysipelotrichia*, *Mucorales*, *Staphylococcus*, and *Mycobacteria*. These bacteria are recognized for their association with various infectious diseases in humans ^41–44^. Current evidence suggests prior infections as a risk factor and the most common triggers for DKA in children with T1D ^45–47^. The increased abundance of these pathogens suggests a potential gut-origin infection and gut dysbiosis in children with DKA. However, more research is needed to clarify the causal relationships between gut dysbiosis, gut infection, and the development of DKA.

This study has some limitations. Firstly, it is a cross-sectional study revealing gut microbiota and metabolite alterations in DKA children, yet further research is needed to clarify the mechanisms underlying SCFA changes. Secondly, intervention studies with SCFAs or SCFA-producing probiotics are necessary to establish causality between SCFAs and DKA. Nevertheless, our study is the first to identify SCFAs and their produces as potential omics biomarkers of DKA in Chinese children with T1D, shedding light on novel therapeutic strategies. Finally, this study was conducted exclusively with Chinese children; larger multicenter studies with expanded sample sizes are needed to validate the findings.

In summary, this study investigates the interplay between clinical characteristics, gut microbiota, and metabolic products in children with T1D, revealing a significant reduction in SCFA-producing bacteria and fecal SCFA levels during the acute onset of DKA. This alteration is negatively correlated with several key clinical features of DKA. These findings suggest that SCFA-producing bacteria and SCFAs may be protective factors against DKA, potentially acting as probiotics or prebiotics with therapeutic effects during the acute phase of the condition.

## Acknowledgements

This study is funded by the Noncommunicable Chronic Diseases-National Science and Technology Major Project (2023ZD0508200 and 2023ZD0508204), National Key Research and Development Program of China (2021YFC2701900), and Zhejiang Provincial Key Science and Technology Project (LGF21H070004). The authors declare no conflicts of interest associated with this manuscript.

## Data Availability

All raw data and relevant results produced in this study are available from the corresponding author upon reasonable request.

---

**Supplementary Table 1.**
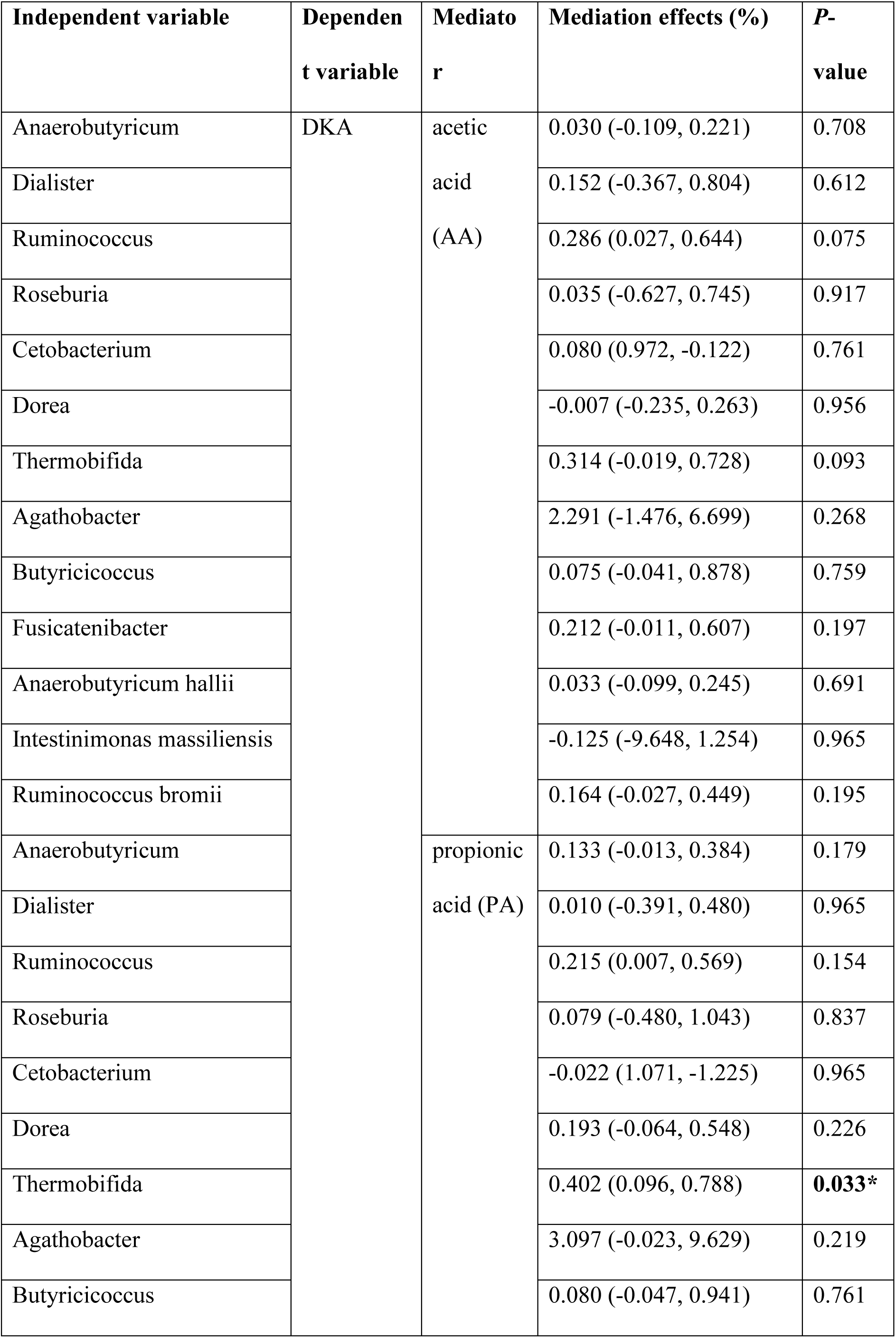
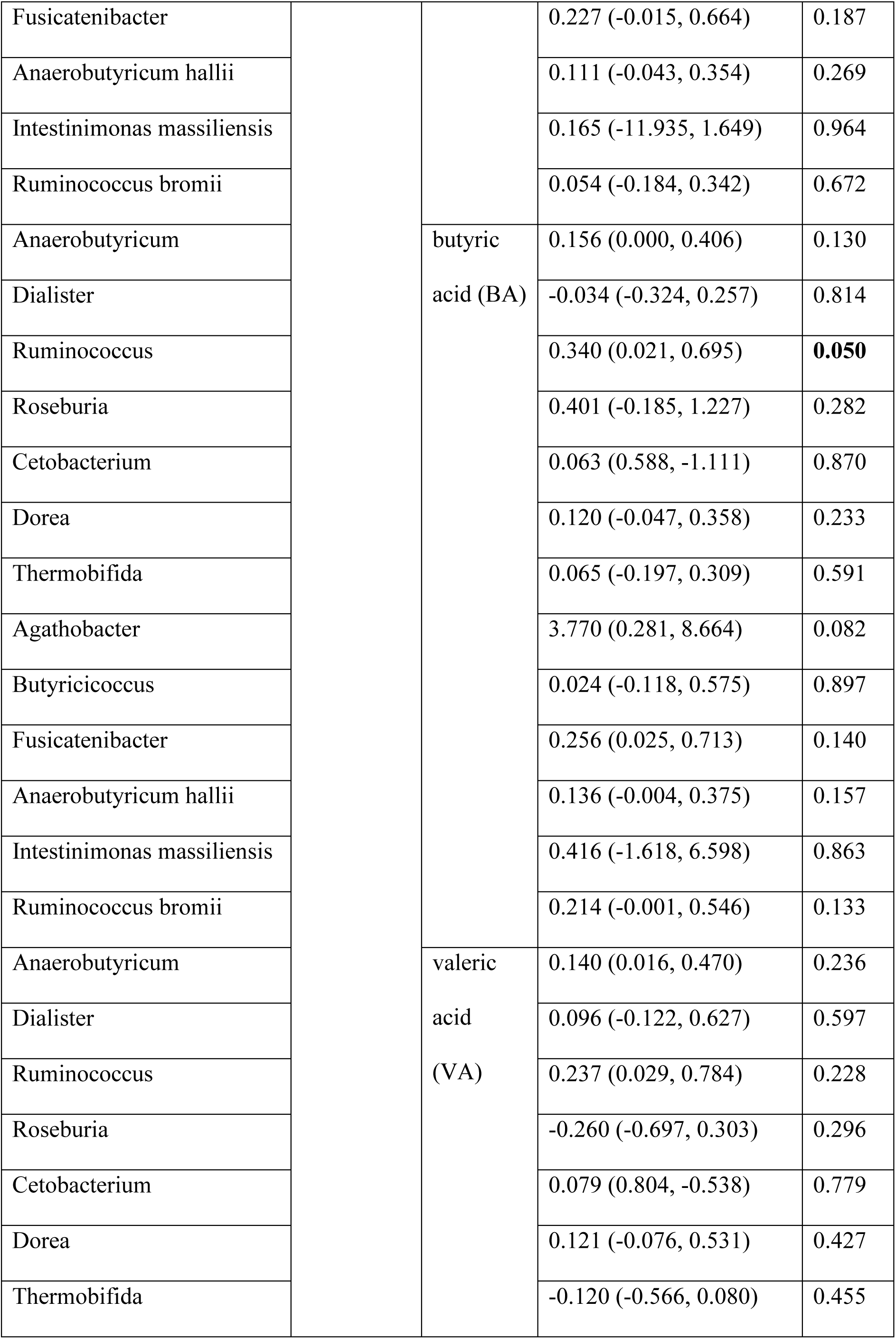
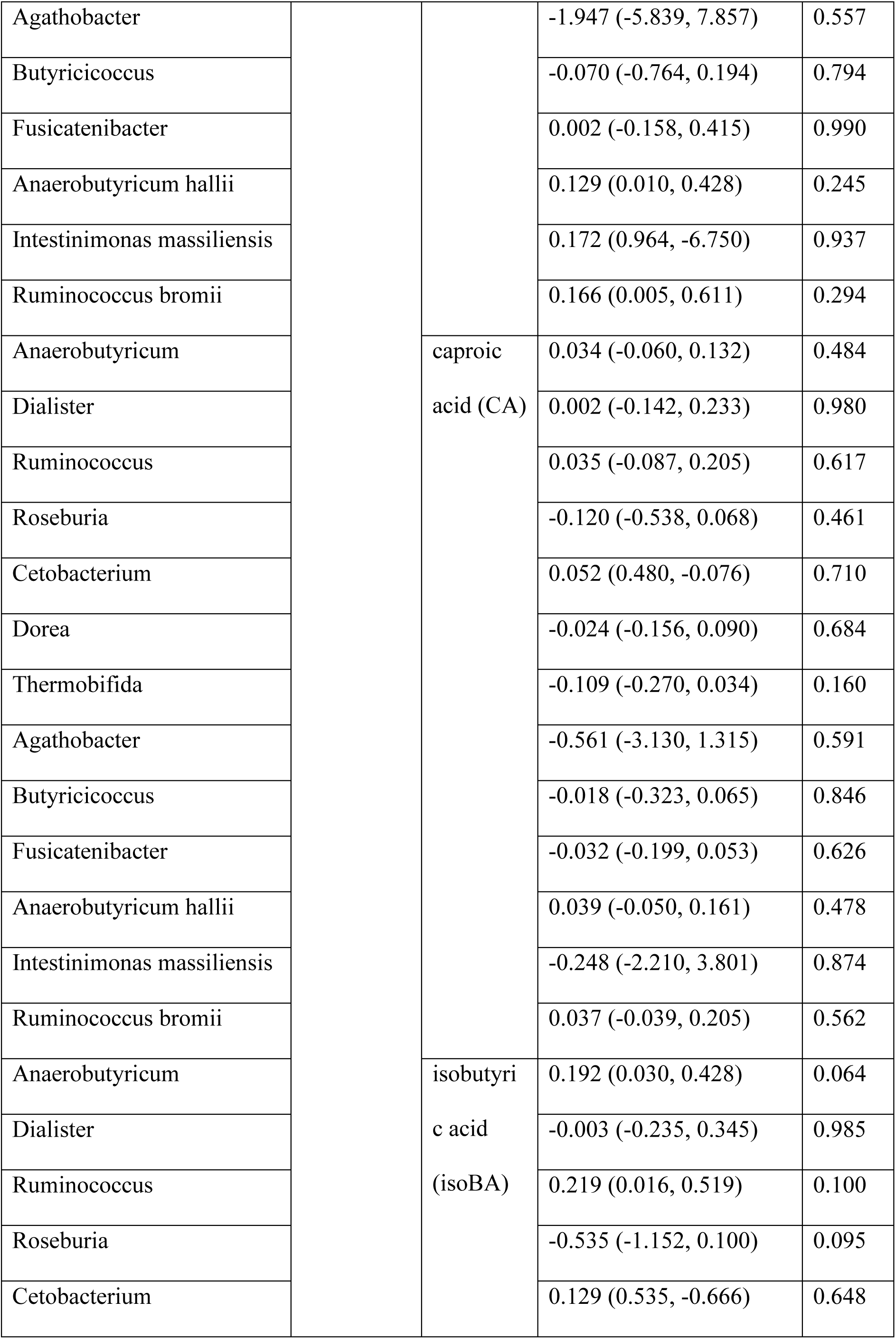
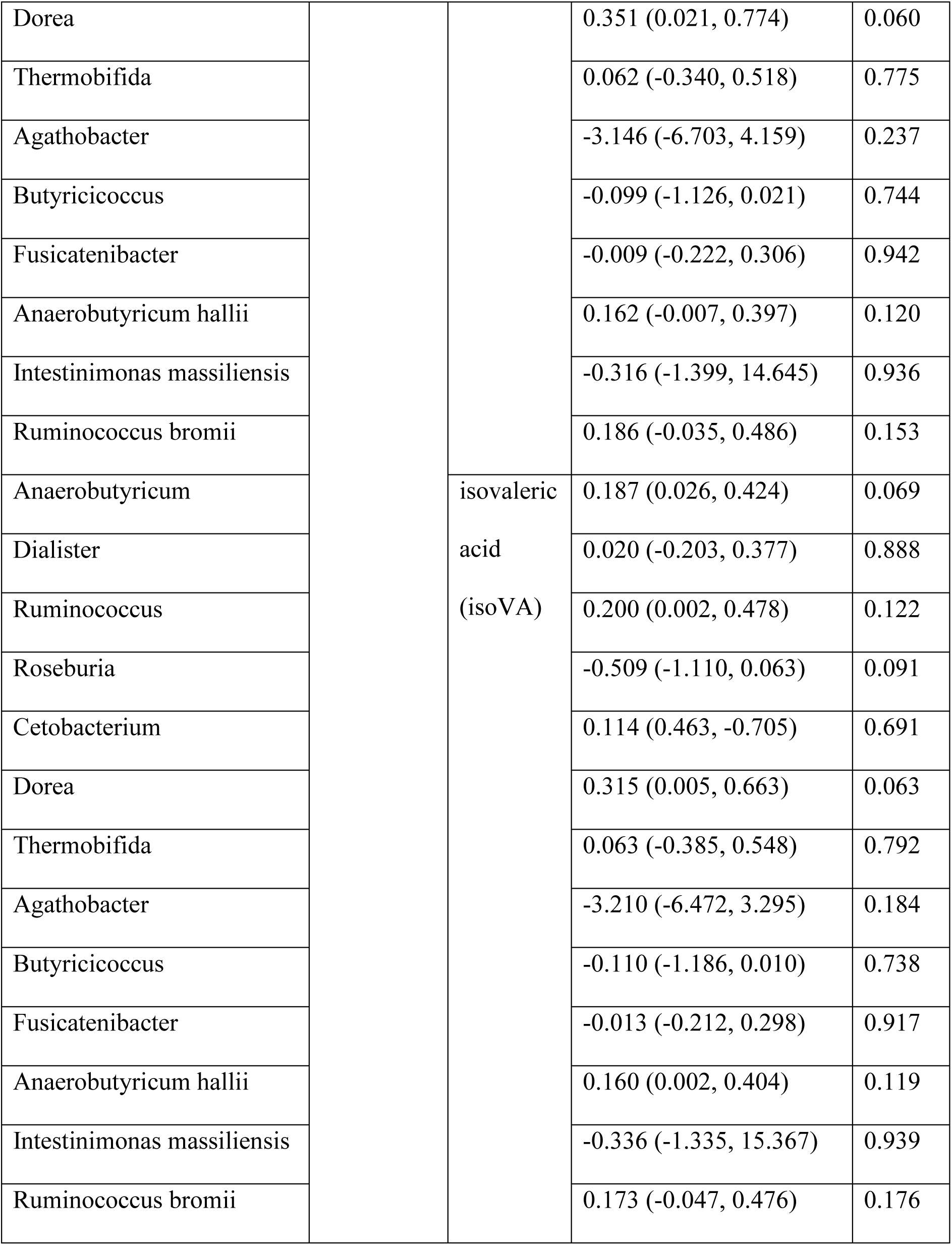
Mediation effects of SCFAs in the association between microbes and DKA. * *P* < 0.05.

**Supplementary Figure 1.**
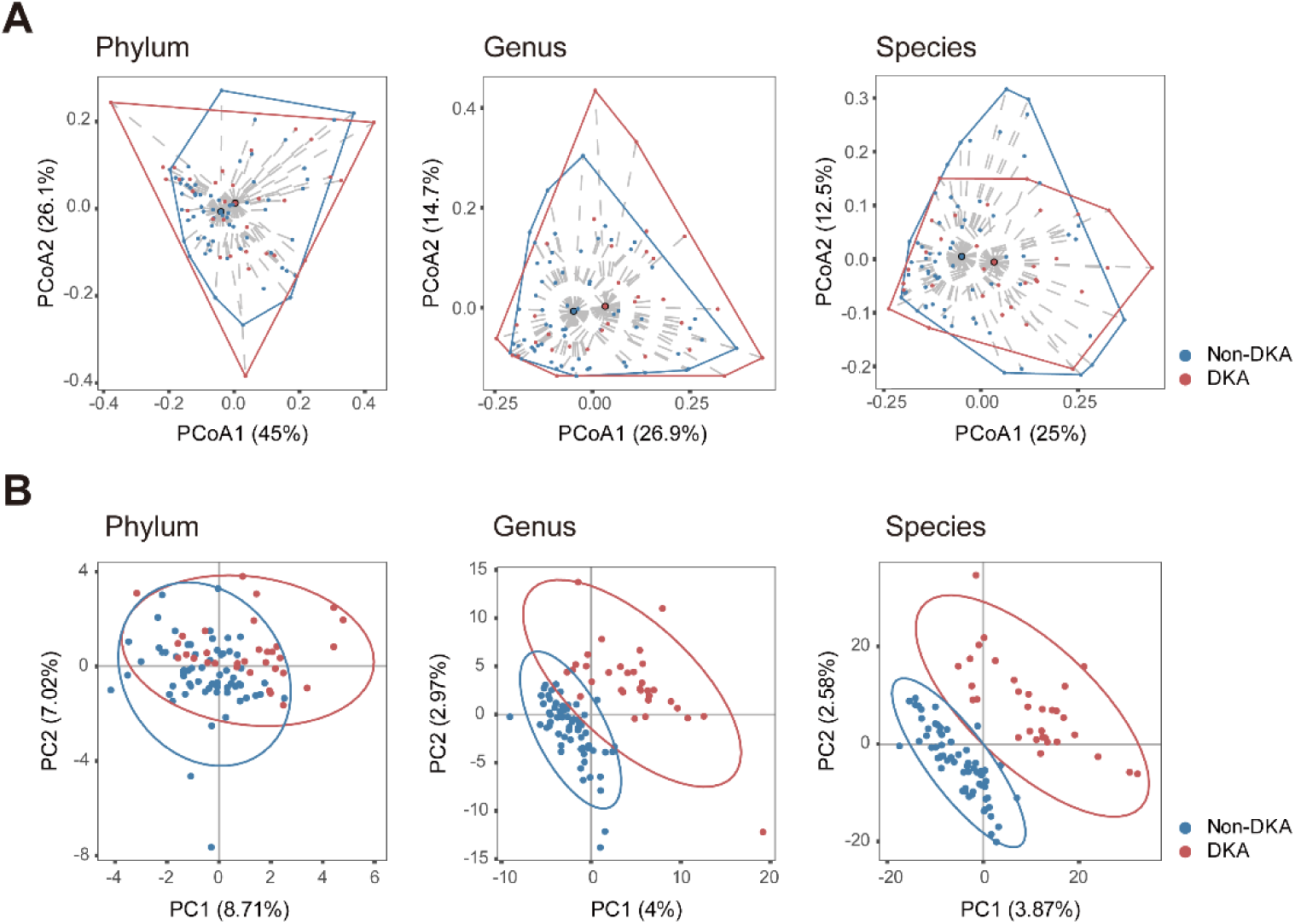
Gut microbiota composition of DKA and non-DKA groups analyzed by PCoA and PLS-DA. (**A**) Gut microbiota beta-diversity of DKA and non-DKA groups differed at the phylum, genus, and species levels through principal coordinates analysis (PCoA). For phylum, *P* = 0.028; for genus, *P* = 0.013; for species, *P* = 0.010. (**B**) Through partial least squares discriminant analysis (PLS-DA), DKA and non-DKA groups were distinguished at the phylum, genus, and species level (all *P* < 0.05).

**Supplementary Figure 2.**
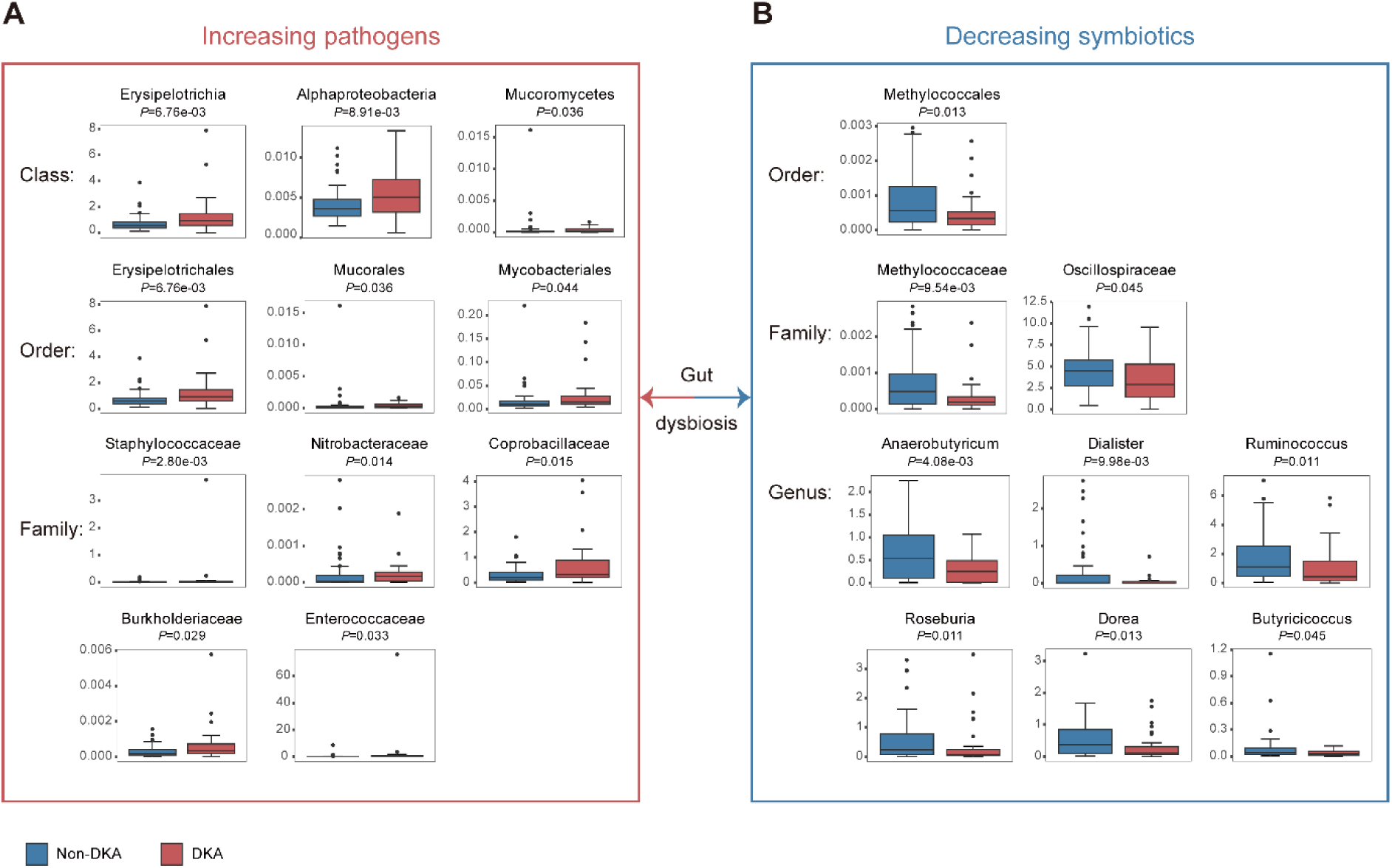
The imbalance of pathogenic and symbiotic microbiota in the DKA group. (**A**) Potential pathogens were significantly increased in the DKA group at the class, order, and family levels. (**B**) Symbiotics with potential beneficial effects in the host were less abundant in the DKA group compared to the non-DKA group.

## Abbreviations

T1D: type 1 diabetes;
DKA: diabetic ketoacidosis;
SCFAs: short-chain fatty acids;
Treg: regulatory T;
TEDDY: the Environmental Determinants of Diabetes in the Young;
FMT: fecal microbiota transplantation;
BMI: body mass index;
HbA1c: glycated hemoglobin;
PCoA: principal coordinates analysis;
PLS-DA: partial least squares discriminant analysis;
AA: acetic acid, or acetate;
PA: propionic acid, or propionate;
BA: butyric acid, or butyrate;
VA: valeric acid, or valerate;
CA: caproic acid, or hexanoate;
IsoBA: isobutyric acid;
IsoVA: isovaleric acid;
CI: confidence interval;
GC/MS: gas chromatography/mass spectrometry;
SBP: systolic blood pressure;
DBP: diastolic blood pressure;
ABE: actual base excess;
HDL-c: high-density lipoprotein cholesterol;
LDL-c: low-density lipoprotein cholesterol;
TG: triglycerides;
ALT: alanine transaminase;
AST: aspartate aminotransferase;
WBC: white blood cells;
TSH: thyroid-stimulating hormone;
T3: triiodothyronine;
T4: thyroxine;
fT3: free triiodothyronine;
fT4: free thyroxine;
IGF1: insulin-like growth factor 1;
IGFBP3: insulin-like growth factor binding protein 3;
RBP: retinol-binding protein;
C3: complement component 3;
IL10: interleukin 10;
CD3: CD3 positive T cells;
CD4: CD4 positive T cells;
CD8: CD8 positive T cells;
Urine pr/cr: urine protein to creatinine;
IDAA1c: insulin dose-adjusted glycated hemoglobin A1c;
NTI: non-thyroidal illness syndrome;
ESS: euthyroid sick syndrome;
IE: indirect effect;
DE: direct effect;
GPR43: G-protein coupled receptor 43;
PYY: peptide YY;
GLP-1: glucagon-like peptide-1.

